# Impaired cholesterol metabolism in the mice model of cystic fibrosis. A vicious circle?

**DOI:** 10.1101/2019.12.13.875336

**Authors:** Felice Amato, Alice Castaldo, Giuseppe Castaldo, Gustavo Cernera, Gaetano Corso, Eleonora Ferrari, Monica Gelzo, Romina Monzani, Valeria Rachela Villella, Valeria Raia

**Author notes:** All the authors equally participated in the study, coordinated by Prof. Luigi Maiuri who is no longer with us. Luigi, with his enthusiasm and his great scientific competence, guided us in this and in many other studies that we conducted with him.

## Abstract

Patients with cystic fibrosis (CF) have low cholesterol absorption and, despite enhanced endogenous biosynthesis, low serum cholesterol. Herein, we investigated cholesterol metabolism in a murine CF model in comparison to *wild type* (WT) testing serum and liver surrogate biomarkers together with the hepatic expression of genes involved in cholesterol metabolism. CF mice display lower sterols absorption and increased endogenous biosynthesis. Subsequently, we evaluated the effects of a cholesterol-supplemented diet on cholesterol metabolism in CF and WT mice. The supplementation in WT mice determines biochemical changes similar to humans. Instead, CF mice with supplementation did not show significant changes, except for serum phytosterols (−50%), liver cholesterol (+35%) and TNFα mRNA expression, that resulted 5-fold higher than in CF without supplementation. However, liver cholesterol in CF mice with supplementation resulted significantly lower compared to WT supplemented mice. This study shows that in CF mice there is a vicious circle in which the altered bile salts synthesis/secretion contribute to reduce cholesterol digestion/absorption. The consequence is the enhanced liver cholesterol biosynthesis that accumulates in the cell triggering inflammation.

## Introduction

Cystic fibrosis (CF) is a life-limiting, autosomal recessive genetic disorder caused by mutations in the *cystic fibrosis transmembrane conductance regulator* (*CFTR*) gene. These mutations lead to a defective transport of chloride and other ions through the respiratory, biliary, gastrointestinal and reproductive epithelia bringing on the secretion of thick mucus (Cantin *et al*, 2015). In particular, at pancreatic level, the mucus represents an obstacle to the secretion of pancreatic enzymes in the intestine causing pancreatic insufficiency (PI) with an altered digestion (Gibson-Corley *et al*, 2016; Peretti *et al*, 2005) and absorption of lipids (Turck *et al*, 2016).

In a preliminary study (Gelzo *et al*, 2016), we evaluated cholesterol metabolism in patients with CF by analyzing surrogate biomarkers (Miettinen *et al*, 2011). Specifically, we determined the plasma levels of campesterol and β-sitosterol (phytosterols) that are biomarkers of intestinal lipid absorption and lathosterol, as a marker of hepatic *de novo* cholesterol biosynthesis. The study showed reduced intestinal absorption of sterols together with an increase of endogenous cholesterol biosynthesis in patients with CF compared to unaffected subjects. But, despite the increased liver synthesis, plasma cholesterol levels in patients with CF were lower (Gelzo *et al*, 2016). This may be due to the partial retention of endogenous cholesterol in hepatocytes, due to a defective exocytosis of cholesterol in patients with CF, as previously demonstrated in CF cell lines and in animal tissues (White *et al*, 2007).

Thus, we observed that CF patients with PI had a significantly reduced absorption of cholesterol and vitamin E as compared to CF patients with pancreatic sufficiency (PS) and to healthy controls suggesting that the supplementation of pancreatic enzymes used in all CF patients with PI was not sufficient to permit a normal digestion of lipids (unpublished results). This could depend on the small amount of cholesterol esterase in the supplementation therapy (Walters *et al*, 2001) and on the reduced synthesis, secretion (Russell, 2003) and reabsorption of bile salts (van de Peppel *et al*, 2019) that occur in patients with CF.

Therefore, the aims of the present study were: i) to investigate cholesterol metabolism in CF compared to *wild type* (WT) mice by determining serum and liver levels of surrogate biomarkers together with the hepatic expression of genes involved in biosynthesis and catabolism of cholesterol; ii) to evaluate the effects of cholesterol supplementation on cholesterol metabolism in CF and WT mice.

## Results

### *Wild type* mice fed with and without cholesterol-supplemented diet

We compared 4 WT mice that received the cholesterol-supplemented diet with 3 WT mice that received a diet with no cholesterol supplementation. As shown in Table 1A, supplemented mice show significantly lower levels of serum phytosterols (7.6 fold) and liver lathosterol (7.6 fold) and significantly higher levels of liver cholesterol (6.2 fold) and cholestanol (4.2 fold). Serum cholesterol was significantly higher (1.3 fold). Furthermore, as shown in Table 1B, WT mice fed with cholesterol supplementation have significantly lower levels of liver LDLR (2.5 fold) and HMG-CoAR (2.2 fold) mRNA, while ACAT2 and CYP7A1 mRNA levels resulted significantly higher (2.9 and 6.0 fold, respectively). Finally, liver mRNA levels of TNFα were higher although not significantly (Figure 1).

**Table 1A.** Comparison of serum and liver sterols in *wild type* (WT) mice fed with and without cholesterol-supplemented diet.

| Mouse populations | Serum |  | Liver |  |  |
| --- | --- | --- | --- | --- | --- |
|  | Phytosterols<br>(mg/dL) | Cholesterol<br>(mg/dL) | Lathosterol<br>(ng/mg tissue) | Cholesterol<br>(μg/mg tissue) | Cholestanol<br>(ng/mg tissue) |
| WT with supplemented diet | 0.25 (0.08) | 122.1 (8.3) | < 0.2 | 15.1 (4.6) | 36.0 (7.7) |
| WT with no supplemented diet | 1.92 (0.37) | 89.9 (2.5) | 1.5 (0.1) | 2.1 (0.2) | 8.7 (1.7) |
| p level of significance | < 0.01 | < 0.001 | < 0.001 | < 0.01 | < 0.001 |

**Table 1B.** Comparison of relative liver expression of LDLR, HMG-CoAR, ACAT2 and CYP7A1 mRNA in *wild type* (WT) mice fed with and without cholesterol-supplemented diet.

| Mouse populations | LDLR | HMG-CoAR | ACAT2 | CYP7A1 |
| --- | --- | --- | --- | --- |
| WT with supplemented diet | 1.0 (0.3) | 5.3 (1.0) | 22.7 (3.7) | 11.8 (6.1) |
| WT with no supplemented diet | 2.5 (0.5) | 11.7 (3.7) | 7.8 (0.8) | 2.0 (1.5) |
| p level of significance | < 0.005 | < 0.02 | < 0.002 | < 0.05 |

**Figure 1.**
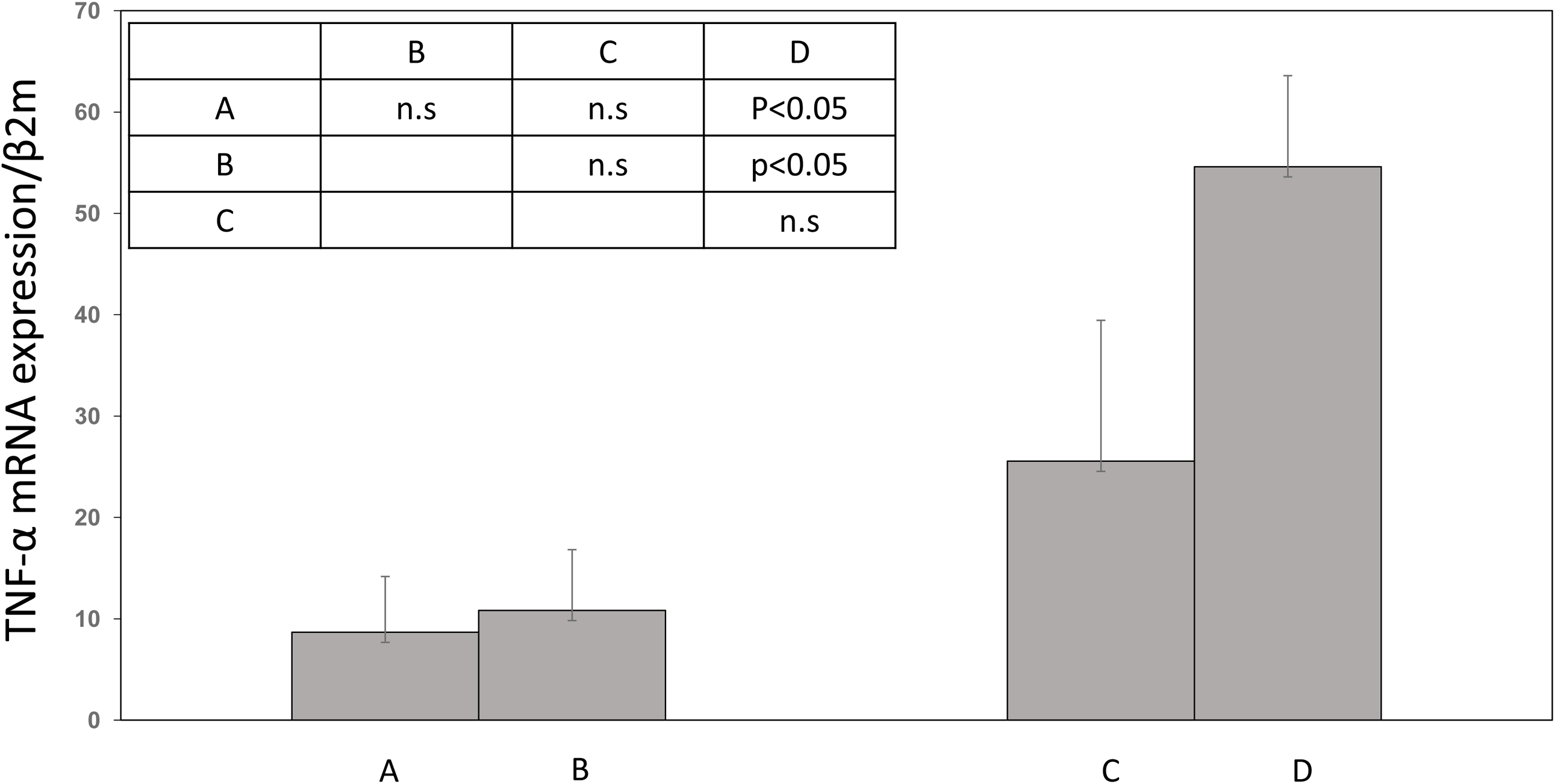
Relative liver expression of TNFα mRNA in mice normalized on expression levels of β-2–microglobulin (β2m). A) *Wild type* (WT) mice fed without cholesterol-supplemented diet. B) CF mice fed without cholesterol-supplemented diet. C) WT mice fed with cholesterol-supplemented diet. D) CF mice fed with cholesterol-supplemented diet. Error bars correspond to standard error.

### CF and WT mice fed without cholesterol-supplemented diet

We compared 3 CF mice with 3 WT mice both fed with no cholesterol-supplemented diet. As shown in Table 2A, CF mice have significantly lower levels of serum phytosterols (2.0 fold), significantly higher levels of liver lathosterol (3.1 fold) and cholesterol (1.2 fold), while liver cholestanol was not significantly different. Serum cholesterol was lower (although not significantly) as compared to WT mice. Furthermore, as shown in Table 2B, CF mice have higher levels of liver LDLR and ACAT2 (although not significantly), while mRNA levels of HMG-CoAR (18.0 fold) and CYP7A1 (16.3 fold) mRNA were significantly higher as compared to WT mice. Finally, liver mRNA levels of TNFα were higher, although not significantly in CF mice (Figure 1).

**Table 2A.** Comparison of serum and liver sterols in cystic fibrosis (CF) and *wild type* (WT) mice both fed with no cholesterol-supplemented diet (n.s.: not significant, p > 0.05).

| Mouse populations | Serum |  | Liver |  |  |
| --- | --- | --- | --- | --- | --- |
| | Phytosterols<br>(mg/dL) | Cholesterol<br>(mg/dL) | Lathosterol<br>(ng/mg tissue) | Cholesterol<br>(μg/mg tissue) | Cholestanol<br>(ng/mg tissue) |
| CF with no supplemented diet | 0.97 (0.14) | 81.7 (6.3) | 4.7 (0.7) | 2.6 (0.1) | 9.5 (1.1) |
| WT with no supplemented diet | 1.92 (0.37) | 89.9 (2.5) | 1.5 (0.1) | 2.1 (0.2) | 8.7 (1.7) |
| p level of significance | < 0.01 | n.s. | < 0.01 | < 0.02 | n.s. |

**Table 2B.** Comparison of relative liver expression of LDLR, HMG-CoAR, ACAT2 and CYP7A1 mRNA in CF mice and *wild type* (WT mice), both fed with no cholesterol-supplemented diet (n.s.: not significant, p > 0.05).

| Mouse populations | LDLR | HMG-CoAR | ACAT2 | CYP7A1 |
| --- | --- | --- | --- | --- |
| CF with no supplemented diet | 5.7 (2.4) | 207.0 (52.1) | 18.4 (6.2) | 32.7 (4.8) |
| WT with no supplemented diet | 2.5 (0.5) | 11.7 (3.7) | 7.8 (0.8) | 2.0 (1.5) |
| p level of significance | n.s. | < 0.05 | n.s. | < 0.0005 |

### CF mice fed with cholesterol-supplemented diet

We compared 4 CF mice that received a diet with cholesterol supplementation with either the 4 WT mice that received a diet with cholesterol supplementation and with the 3 CF mice that received a diet with no cholesterol supplementation. As shown in Table 3A, comparing the CF with WT mice both with cholesterol supplemented diet, we observed: i) significantly higher levels of serum phytosterols (2.0 folds) and liver lathosterol (22.0 fold); ii) significantly lower levels of both serum (1.3 fold) and liver cholesterol (4.3 fold) together with significantly lower liver cholestanol (3.4 fold). Furthermore, as shown in Table 3B, CF mice fed with cholesterol supplementation have significantly higher levels of liver LDLR (4.7 fold), HMG-CoAR (41.2 fold) and CYP7A1 (5.3 fold) mRNA levels as compared to WT mice fed with cholesterol supplementation, while the liver mRNA levels of ACAT2 were not significantly different (Table 3B). Finally, liver mRNA levels of TNFα were higher, although not significantly (Figure 1).

**Table 3A.**
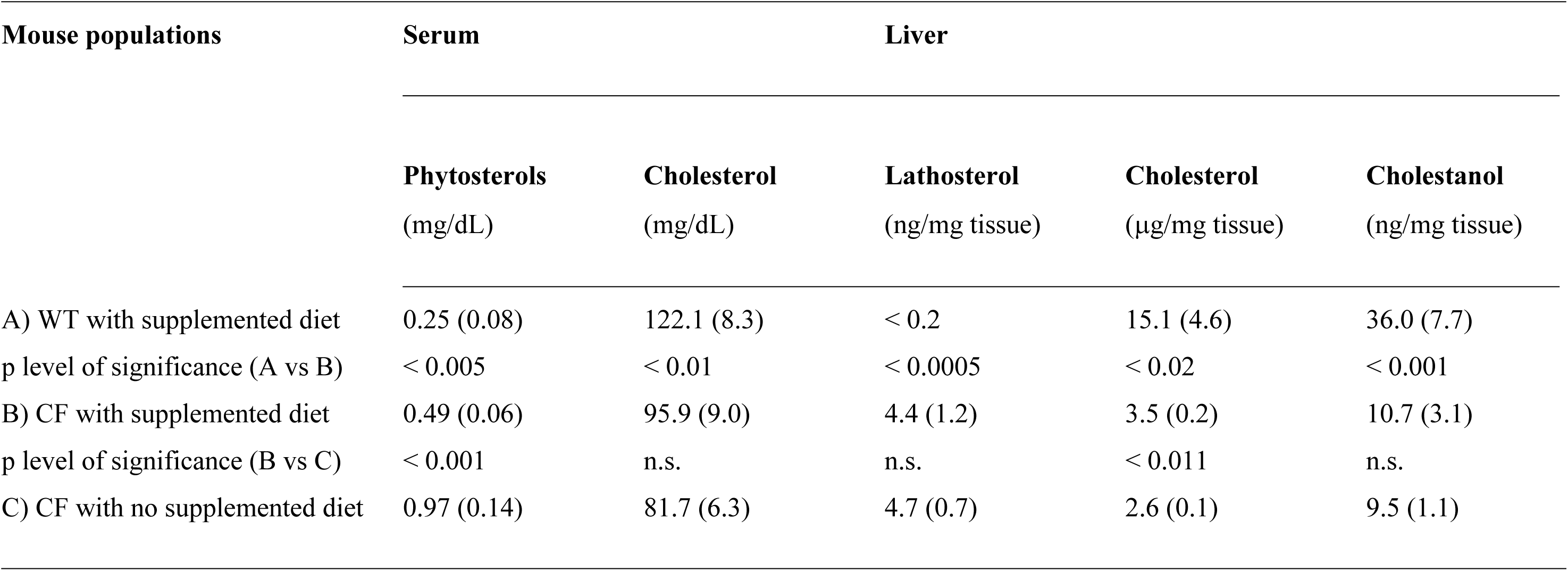
Serum and liver sterols in CF mice fed with cholesterol-supplemented diet in comparison to either *wild type* (WT) mice fed with cholesterol-supplemented diet and CF mice fed with no cholesterol-supplemented diet (n.s.: not significant, p > 0.05).

**Table 3B.** Relative liver expression of LDLR, HMG-CoAR, ACAT2 and CYP7A1 mRNA in CF mice fed with cholesterol-supplemented diet in comparison to either *wild type* (WT) mice fed with cholesterol-supplemented diet and CF mice fed with no cholesterol-supplemented diet (n.s.: not significant, p > 0.05).

| Mouse populations | LDLR | HMG-CoAR | ACAT2 | CYP7A1 |
| --- | --- | --- | --- | --- |
| A) WT with supplemented diet | 1.0 (0.3) | 5.3 (1.0) | 22.7 (3.7) | 11.8 (6.1) |
| p level of significance (A vs B) | n.s. | < 0.02 | n.s. | < 0.05 |
| B) CF with supplemented diet | 4.7 (3.0) | 206.0 (38.1) | 29.3 (18.4) | 63.2 (40.0) |
| p level of significance (B vs C) | n.s. | n.s. | n.s. | n.s. |
| C) CF with no supplemented diet | 5.7 (2.4) | 207.0 (52.1) | 18.4 (6.2) | 32.7 (4.8) |

Finally, as shown in Table 3A and B, comparing the CF mice with supplemented diet to CF mice with no supplemented diet, we observed that all the parameters resulted not significantly different, with the exception of serum phytosterols that were significantly lower in CF mice supplemented (2.0 fold), liver cholesterol (1.3 fold) and liver mRNA levels of TNFα (5.0 fold) that were significantly higher (Figure 1).

## Discussion

Our study demonstrates that the cholesterol supplementation in WT mice results in a series of biochemical changes similar to those physiologically observed in humans (Hu *et al*, 2010), in agreement with previous studies (Repa *et al*, 2004; Schwarz *et al*, 2001). In fact, the supplementation causes: i) enhanced intestinal absorption of cholesterol; ii) reduction of serum phytosterol for a competitive mechanism of intestinal absorption with cholesterol (Gylling & Simonen, 2015); iii) the inhibition of LDLR expression due to the increased liver uptake of cholesterol (Hu *et al*, 2010; Goldstein & Brown, 1990); iv) the inhibition of endogenous biosynthesis of cholesterol, supported by lower liver expression of HMG-CoAR (Hu *et al*, 2010; Goldstein & Brown, 1990) and the consequent decreased liver levels of lathosterol; and v) the activation of liver cholesterol catabolism supported by the enhanced expression of ACAT2 and CYP7A1 that stimulate the esterification of cholesterol and the synthesis of bile salts respectively, and increased liver cholestanol, a catabolic product of acidic pathway of bile salts (Russell, 2003).

Cystic fibrosis mice compared to WT, both fed with no cholesterol-supplemented diet, display a series of metabolic alterations some of which are similar to those observed in patients with CF (Gelzo *et al*, 2016): i) lower intestinal absorption of sterols, supported by the lower levels of serum phytosterols; ii) increased endogenous biosynthesis of cholesterol, confirmed by a very high expression of liver HMG-CoAR and by higher concentrations of liver lathosterol. Nevertheless, the enhanced synthesis of cholesterol is unable to correct the lower levels of serum cholesterol observed in CF as compared to WT mice. These data confirm that in CF mice the liver mechanisms of synthesis/excretion of cholesterol are impaired, as it has been demonstrated in CF cell models and in tissues from CF mice (Fang *et al*, 2010; White *et al*, 2004). That study reported the enhanced *de novo* synthesis of cholesterol, followed by its accumulation at endo-lysosomal level due to a block in the translocation to the Golgi and endoplasmic reticulum (ER). They postulated that the accumulation was due to a generalized mechanism caused by misfolded membrane proteins that escape ER quality control and impact on lipid homeostasis (Gentzsch *et al*, 2007). The lack of cholesterol provision to the ER is followed by the activation of sterol regulatory element-binding protein that enhances the endogenous synthesis of cholesterol (White *et al*, 2007). We confirm that in CF mice liver there is an enhanced biosynthesis of endogenous cholesterol due to the lack of inhibition of HMGCoAR (Hu *et al*, 2010; Goldstein & Brown, 1990), which could be due to the lack of unidentified endogenous factors and causes the accumulation of cholesterol in the liver. Despite this, we observed a lack of inhibition of LDLR expression in the CF liver that further contributes to accumulate cholesterol. Such accumulation triggers inflammation, as it is demonstrated by the higher levels of TNFα expression (Yang & Seki, 2015) that we observed in CF as compared to the WT mice liver. Interestingly, the liver of CF mice enhances the expression of ACAT2, the enzyme that esterifies cholesterol before its secretion (Hu *et al*, 2010; Goldstein & Brown, 1990), but this activation is not followed by a significant release of cholesterol in blood. Similarly, the liver of CF mice expresses very high levels of CYP7A1, the key enzyme in the catabolism of cholesterol to bile salt (Russell, 2003), but also such metabolism may be impaired, since the levels of liver cholestanol, an intermediate of bile salt synthesis (Russell, 2003), are similar to those observed in WT mice.

The cholesterol supplementation in CF mice causes an increase of serum cholesterol (even if less significant of that observed in supplemented WT) and a significant reduction of serum phytosterols (about 50% less) as compared to CF mice with no supplementation, indicating that the supplementation favors some absorption of cholesterol in CF mice. However, the supplementation promotes a mild increase (+ 35%) of cholesterol in the liver, compared to CF mice with no supplementation, which is due to the alterations above mentioned observed in the CF liver increasing further the inflammatory reaction, supported by the considerable expression of TNFα.

All these results show that in CF mice there is a vicious circle, in which the altered synthesis and secretion of biliary salts contribute to reduce cholesterol digestion and absorption; as a consequence there is the enhanced liver biosynthesis of cholesterol that accumulates in the cell triggering inflammation that involves small bile ducts further impairing the synthesis and release of biliary salts. These results have a translational impact on cholesterol metabolism in patients with CF that mostly show low levels of serum cholesterol with a myriad of clinical consequences (Segoviano-Mendoza *et al*, 2018; Elam, 1995). In order to break the vicious circle of CF cholesterol metabolism it would be necessary to act in two directions: i) enhance the cholesterol supplementation with the diet, associated to an adequate amount of bile salts and pancreatic cholesterol esterase for its digestion; ii) modulate the endogenous synthesis in order to reduce cholesterol accumulation in the liver and the consequent inflammation. In this context, the analysis of surrogate markers of cholesterol metabolism may help to define and monitor the individual therapy in patients with CF.

## Materials and methods

### Mice and treatments

Cystic fibrosis mice homozygous for the F508del *CFTR* mutation in the FVB/129 outbred background (Cftrtm1EUR, F508del, FVB/129) and WT littermates, male and female, were obtained from Bob Scholte, Erasmus Medical Center Rotterdam, The Netherlands: CF coordinated action program EU FP6 LSHMCT-2005-018932 (vanDoornick *et al*, 1995).

A first group of 6 week old, CF (n. 3) and WT (n. 3) mice were fed with CF diet (Charles River V1124-70) enriched of linoleic acid and vitamin E for 14 weeks. A second group of 6 week old CF (n. 4) and WT (n. 4) mice, were fed with CF diet (Charles River V1124-70) enriched of linoleic acid and vitamin E for 10 weeks and then they were fed with a diet supplemented of cholesterol (1% wt/wt) for 4 weeks (Envigo TD02026). At the end of the last daily treatment, mice were anesthetized with Avertin (tribromoethanol, 250 mg/kg, Sigma-Aldrich, T48402) and then killed. Blood and liver samples were collected and stored at −80 °C (vanDoornick *et al*, 1995).

All the procedures in mice were approved by the local Ethics Committee for Animal Welfare (IACUC No. 849) and were carried out in strict respect of European and National regulations for animal use in research (2010/63 UE).

### Sterol profile analysis

For the analysis of serum sterols, blood samples without anticoagulant/additive were collected and stored in plastic tubes at −80°C suddenly after collection. Before sterols extraction, each blood sample was centrifuged at 14000 rpm for 10 min at 4°C and 100 μL of supernatant (hemolyzed serum) was collected and placed in a pyrex tube. To precipitate the hemoglobin, 1.7 mL of cool ethanol was added to the sample, vigorously mixed and incubated in ice for 15 min. The sample was then centrifuged at 3000 rpm for 25 min and the supernatant was transferred in a second pyrex tube for the sterols extraction performed as previously described (Gelzo *et al*, 2016; Corso *et al*, 2011). Briefly, the sample was mixed with 40 μg of 5α-cholestane (internal standard) and hydrolyzed by incubation in 1N potassium hydroxide ethanolic solution at 80°C for 1 hour. The sterols were then extracted by hexane and derivatized by BSTFA and pyridine, dried under nitrogen, and dissolved in 100 μL of dichloromethane. For qualitative sterols analysis, 1 μL of solution was injected in a gas chromatograph coupled with a mass spectrometer (GC-MS; GC 7890 A/MD 5975C, Agilent Technologies, Santa Clara, CA, USA) in scan mode from *m/z* 50 to 500. While, quantitative analysis was carried out by injecting 1 μL of solution in a gas chromatograph coupled with a flame ionization detector (GC-FID; HP-5890, Agilent Technologies). Both instrumentations were equipped with Elite-5MS capillary column (PerkinElmer, USA).

For the analysis of sterols in the liver, the tissue samples were first weighed and the homogenates were subsequently prepared as follows. The liver tissue was placed in a clean plastic vial with 400-800 μL of distilled water, based on the weight of the tissue, and homogenized using an Ultra Turrax T25 digital homogenizer (IKA®-Werke GmbH & Co. KG, Staufen, Germany). Sterols were extracted from 100 μL of liver homogenate and analyzed as described above for serum. The levels of sterols in liver were normalized by tissue weight and expressed as μg/mg of tissue for cholesterol and ng/mg of tissue for other sterols. The analytical solvents of HPLC grade were obtained from Sigma (St Louis, MO, USA). Potassium hydroxide was purchased from Merck (Merck KGaA, Darmstadt, Germany). N,O-bis(trimethylsilyl)trifluoroacetamide (BSTFA) was obtained from Sigma (St Louis, MO, USA). Stock solutions of standard sterols (Sigma, St Louis, MO, USA) were prepared in chloroform/methanol (2:1, v/v) at concentration of 1 mg/mL. For 5α-cholestane (internal standard for sterol analysis), a work solution at concentration of 0.4 mg/mL was prepared.

### RNA isolation, qRT-PCR

We analyzed by qRT-PCR the levels of expression of the following genes: *low density lipoprotein receptor (LDLR), 3-hydroxy-3-methylglutaryl-CoA reductase (HMG-CoAR), acyl CoA:cholesterol acyl transferase 2 (ACAT2), cytochrome P450 7A1 (CYP7A1), tumor necrosis factor alpha (TNFα).*

Total RNA was extracted from mouse liver tissue using TriZol (Invitrogen, Waltham, MA, USA) according to the manufacturer’s instructions and genomic DNA was removed by treating with DNAse enzyme. RNA concentration and purity were measured using NanoDrop 2000 (Thermo Scientific). RNA (1 μg) was converted to cDNA by reverse transcription using iScript cDNA synthesis kit (Bio-Rad).

qRT-PCR was performed using iQ SYBR Green Supermix (Bio-Rad) and the ABI 7900 HT Real Time PCR System (Applied Biosystems). The qRT-PCR results were normalized on expression levels of β-2–microglobulin, used as internal reference.

### Statistical analysis

Data were reported as mean and standard deviation. Comparisons between two groups were performed by T-test. The significance was accepted at the level of p < 0.05.

## Non-standard abbreviations

ACAT2: acyl CoA:cholesterol acyl transferase 2;
CF: cystic fibrosis;
CFTR: CF transmembrane conductance regulator;
CYP7A1: cytochrome P450 7A1;
ER: endoplasmic reticulum;
HMG-CoAR: 3-hydroxy-3-methylglutaryl-CoA reductase;
LDLR: low density lipoprotein receptor;
PI: pancreatic insufficiency;
PS: pancreatic sufficiency;
TNFα: tumor necrosis factor alpha;
WT: wild type.

## Acknowledgements

We gratefully acknowledge Regione Campania (Quota vincolata per la prevenzione e cura della Fibrosi Cistica L. 548/94, Ricerca). FSN 2015, 2016, 2017 and 2018.

## Author contributions

All the authors equally participated in the study.

## Conflict of interest

The authors declare that they have no conflict of interest.

## Data Availability Section

The primers used for qRT-PCR are available on request.

## References

Cantin AM, Hartl D, Konstan MW, Chmiel JF (2015) Inflammation in cystic fibrosis lung disease: pathogenesis and therapy. J Cyst Fibros 14: 419–430

Corso G, Gelzo M, Barone R, Clericuzio S, Pianese P, Nappi A, Dello Russo A (2011) Sterol profiles in plasma and erythrocyte membranes in patients with Smith-Lemli-Opitz syndrome: a six-year experience. Clin Chem Lab Med 49: 2039–2046

Elam B (1995) Increased mortality in older persons with hypocholesterolemia: cause or effect. J Am Geriatr Soc 43: 312–313

Fang D, West RH, Manson ME, Ruddy J, Jiang D, Previs SF, Sonawane ND, Burgess JD, Kelley TJ (2010) Increased plasma membrane cholesterol in cystic fibrosis cells correlates with CFTR genotype and depends on de novo cholesterol synthesis. Respir Res 11: 61

Gelzo M, Sica C, Elce A, Dello Russo A, Iacotucci P, Carnovale V, Raia V, Salvatore D, Corso G, Castaldo G (2016) Reduced absorption and enhanced synthesis of cholesterol in patients with cystic fibrosis: a preliminary study of plasma sterols. Clin Chem Lab Med 54: 1461–1466

Gentzsch M, Choudhury A, Chang XB, Pagano RE, Riordan JR (2007) Misassembled mutant DeltaF508 CFTR in the distal secretory pathway alters cellular lipid trafficking. J Cell Sci 120: 447–455

Gibson-Corley KN, Meyerholz DK, Engelhardt JF (2016) Pancreatic pathophysiology in cystic fibrosis. J Pathol 238: 311–320

Goldstein JL, Brown MS (1990) Regulation of the mevalonate pathway. Nature 343: 425–430

Gylling H, Simonen P (2015) Phytosterols, Phytostanols, and Lipoprotein Metabolism. Nutrients 7: 7965–7977

Hu YW, Zheng L, Wang Q (2010) Regulation of cholesterol homeostasis by liver X receptors. Clin Chim Acta 411: 617–625

Miettinen TA, Gylling H, Nissinen MJ (2011) The role of serum noncholesterol sterols as surrogate markers of absolute cholesterol synthesis and absorption. Nutr Metab Cardiovasc Dis 21: 765–769

Peretti N, Marcil V, Drouin E, Levy E (2005) Mechanisms of lipid malabsorption in cystic fibrosis: the impact of essential fatty acids deficiency. Nutr Metab 2: 11

Repa JJ, Buhman KK, Farese RV Jr, Dietschy JM, Turley SD (2004) ACAT2 deficiency limits cholesterol absorption in the cholesterol-fed mouse: impact on hepatic cholesterol homeostasis. Hepatology 40: 1088–1097

Russell DW (2003) The enzymes, regulation, and genetics of bile acid synthesis. Annu Rev Biochem 72: 137–174

Schwarz M, Davis DL, Vick BR, Russell DW (2001) Genetic analysis of cholesterol accumulation in inbred mice. J Lipid Res 42: 1812–1819

Segoviano-Mendoza M, Cárdenas-de la Cruz M, Salas-Pacheco J, Vázquez-Alaniz F, La Llave-León O, Castellanos-Juárez F, Méndez-Hernández J, Barraza-Salas M, Miranda-Morales E, Arias-Carrión O et al (2018) Hypocholesterolemia is an independent risk factor for depression disorder and suicide attempt in Northern Mexican population. BMC Psychiatry 18: 7

Turck D, Braegger CP, Colombo C, Declercq D, Morton A, Pancheva R, Robberecht E, Stern M, Strandvik B, Wolfe S et al (2016) ESPEN-ESPGHAN-ECFS guidelines on nutrition care for infants, children, and adults with cystic fibrosis. Clin Nutr 35: 557–577

van de Peppel IP, Bodewes FAJA, Verkade HJ, Jonker JW (2019) Bile acid homeostasis in gastrointestinal and metabolic complications of cystic fibrosis. J Cyst Fibros 18: 313–320

van Doorninck JH, French PJ, Verbeek E, Peters RH, Morreau H, Bijman J, Scholte BJ (1995) A mouse model for the cystic fibrosis delta F508 mutation. EMBO J 14: 4403–4411

Walters MP, Conway SP (2001) Cholesterol esterase activities in commercial pancreatic enzyme preparations and implications for use in pancreatic insufficient cystic fibrosis. J Clin Pharm Ther 26: 425–431

White NM, Corey DA, Kelley TJ (2004) Mechanistic similarities between cultured cell models of cystic fibrosis and niemann-pick type C. Am J Respir Cell Mol Biol 31: 538–543

White NM, Jiang D, Burgess JD, Bederman IR, Previs SF, Kelley TJ (2007) Altered cholesterol homeostasis in cultured and in vivo models of cystic fibrosis. Am J Physiol Lung Cell Mol Physiol 292: L476–L486

Yang YM, Seki E (2015) TNFα in liver fibrosis. Curr Pathobiol Rep 3: 253–261

